# Genetic architecture of gene expression traits across diverse populations

**DOI:** 10.1101/245761

**Authors:** Lauren S. Mogil, Angela Andaleon, Alexa Badalamenti, Scott P. Dickinson, Xiuqing Guo, Jerome I. Rotter, W. Craig Johnson, Hae Kyung Im, Yongmei Liu, Heather E. Wheeler

## Abstract

For many complex traits, gene regulation is likely to play a crucial mechanistic role. How the genetic architectures of complex traits vary between populations and subsequent effects on genetic prediction are not well understood, in part due to the historical paucity of GWAS in populations of non-European ancestry. We used data from the MESA (Multi-Ethnic Study of Atherosclerosis) cohort to characterize the genetic architecture of gene expression within and between diverse populations. Genotype and monocyte gene expression were available in individuals with African American (AFA, n=233), Hispanic (HIS, n=352), and European (CAU, n=578) ancestry. We performed expression quantitative trait loci (eQTL) mapping in each population and show genetic correlation of gene expression depends on shared ancestry proportions. Using elastic net modeling with cross validation to optimize genotypic predictors of gene expression in each population, we show the genetic architecture of gene expression for most predictable genes is sparse. We found the best predicted gene, *TACSTD2*, was the same across populations with R^2^ *>* 0.86 in each population. However, we identified a subset of genes that are well-predicted in one population, but poorly predicted in another. We show these differences in predictive performance are due to allele frequency differences between populations. Using genotype weights trained in MESA to predict gene expression in independent populations showed that a training set with ancestry similar to the test set is better at predicting gene expression in test populations, demonstrating an urgent need for diverse population sampling in genomics. Our predictive models and performance statistics in diverse cohorts are made publicly available for use in transcriptome mapping methods at https://github.com/WheelerLab/DivPop.

**Author summary:** Most genome-wide association studies (GWAS) have been conducted in populations of European ancestry leading to a disparity in understanding the genetics of complex traits between populations. For many complex traits, gene regulation is critical, given the consistent enrichment of regulatory variants among trait-associated variants. However, it is still unknown how the effects of these key variants differ across populations. We used data from MESA to study the underlying genetic architecture of gene expression by optimizing gene expression prediction within and across diverse populations. The populations with genotype and gene expression data available are from individuals with African American (AFA, n=233), Hispanic (HIS, n=352), and European (CAU, n=578) ancestry. After calculating the prediction performance, we found that there are many genes that were well predicted in one population are poorly predicted in another. We further show that a training set with ancestry similar to the test set resulted in better gene expression predictions, demonstrating the need to incorporate diverse populations in genomic studies. Our gene expression prediction models and performance statistics are publicly available to facilitate future transcriptome mapping studies in diverse populations.

## Introduction

For over a decade, genome-wide association studies (GWAS) have facilitated the discovery of thousands of genetic variants associated with complex traits and new insights into the biology of these traits [1]. Most of these studies involved individuals of primarily European descent, which can lead to disparities when attempting to apply this information across populations [2–4]. Continued increases in GWAS sample sizes and new integrative methods will lead to more clinically relevant and applicable results. A recent study shows that the lack of diversity in large GWAS skew the prediction accuracy across non-European populations [5]. This discrepancy in predictive accuracy demonstrates that adding ethnically diverse populations is critical for the success of precision medicine, genetic research, and understanding the biology behind genetic variation [5–8].

Gene regulation is likely to play a critical role for many complex traits as trait-associated variants are enriched in regulatory, not protein-coding, regions [9–13]. Numerous expression quantitative trait loci (eQTLs) studies have provided insight into how genetic variation affects gene expression [14–17]. While eQTLs can act at a great distance, or in *trans*, the largest effect sizes are consistently found near the transcription start sites of genes [14–17]. Because gene expression shows a more sparse genetic architecture than many other complex traits, gene expression is amenable to genetic prediction with relatively modest sample sizes [18, 19]. This has led to new mechanistic methods for gene mapping that integrate transcriptome prediction, including PrediXcan [20] and TWAS [21]. These methods have provided useful tools for understanding the genetics of complex traits; however, most of the models have been built using predominantly European populations.

How the key variants involved in gene regulation differ among populations has not been fully explored. While the vast majority of eQTL mapping studies have been performed in populations of European descent, increasing numbers of transcriptome studies in non-European populations make the necessary comparisons between populations feasible [14, 22, 23]. An eQTL study across eight diverse HapMap populations (*∼* 100 individuals/population) showed that the directions of effect sizes were usually consistent when an eQTL was present in two populations [14]. However, the impact of a particular genetic variant on population gene expression differentiation is also dependent on allele frequencies, which often vary between populations. A better understanding of the degree of transferability of gene expression prediction models across populations is essential for broad application of methods like PrediXcan in the study of the genetic architecture of complex diseases and traits in diverse populations.

Here, in order to better define the genetic architecture of gene expression across populations, we combine genotype [24] and monocyte gene expression [25] data from the Multi-Ethnic Study of Atherosclerosis (MESA) for the first time. We perform eQTL mapping and optimize multi-SNP predictors of gene expression in three diverse populations. The MESA populations studied herein comprise 233 African American (AFA), 352 Hispanic (HIS), and 578 European (CAU) self-reported ancestry individuals. Using elastic net regularization and Bayesian sparse linear mixed modeling, we show sparse models outperform polygenic models in each population. We show the genetic correlation of SNP effects and the predictive performance correlation is highest between populations with the most overlapping admixture proportions. We found a subset of genes that are well predicted in the AFA and/or HIS cohorts that are poorly predicted, if predicted at all, in the CAU cohort. We also test our predictive models trained in MESA cohorts in independent cohorts from the HapMap Project [14], Geuvadis Consortium [26], and Framingham Heart Study [18, 27] and show the correlation between predicted and observed gene expression is highest when the ancestry of the test set is similar to that of the training set. By diversifying our model-building populations, new genes may be implicated in complex trait mapping studies that were not previously interrogated. Models built here have been added to PredictDB for use in PrediXcan [20] and other studies, links at https://github.com/WheelerLab/DivPop.

## Results

### eQTL discovery in MESA and replication in independent populations reflects ancestry and sample size

We surveyed each MESA population (AFA, HIS, CAU) and two combined populations (AFHI, ALL) for cis-eQTLs. SNPs within 1Mb of each of 10,143 genes were tested for association with monocyte gene expression levels using a linear additive model. The MESA HIS cohort includes many individuals with recent African admixture (S1 Fig). We compared models that included a range of genotypic principal components (0, 3, 5,10) and PEER factors (0, 10, 20, 30) to adjust for hidden confounders in the expression data [28]. Genotypic principal components (PCs) and PEER factors were computed within each population prior to cis-eQTL mapping. While 3 genotypic PCs controlled for inflation due to population stratification compared to 0 PCs, especially in HIS, the cis-eQTLs discovered with 3, 5, or 10 PCs were nearly the same (S2 Fig).

We calculated the true positive rates (π_1_) of top cis-eQTLs (FDR *<* 0.05) from our MESA discovery cohorts by examining their P value distribution in several replication cohorts: Framingham Heart Study (FHS, n = 4838 European ancestry individuals, whole blood expression microarray) [27], Geuvadis (GEU, n = 344 European and 77 African ancestry individuals, lympoblastoid cell line (LCL) RNA-Seq) [26], Yoruba from Ibadan, Nigeria (YRI, n = 107, LCL expression microarray) [14], and Mexican ancestry from Los Angeles (MXL, n = 45, LCL expression microarray) [14]. True positive rates were similar across PEER factors except for models with 0 PEER factors, which were either higher or lower depending on the replication population (Fig. 1). Because the FHS replication population is the largest, true positive rates were higher across discovery populations. True positive rates for eQTLs discovered in AFA were higher compared to the other MESA populations in replication populations that include African ancestry individuals (GEU and YRI). eQTLs discovered in AFA and HIS yielded higher true positive rates in both YRI and MXL compared to eQTLs discovered in CAU (Fig. 1). A full pairwise comparison of π_1_statistics across all discovery and replication population PEER factor combinations showed similar trends (S3 Fig).

**Fig 1.**
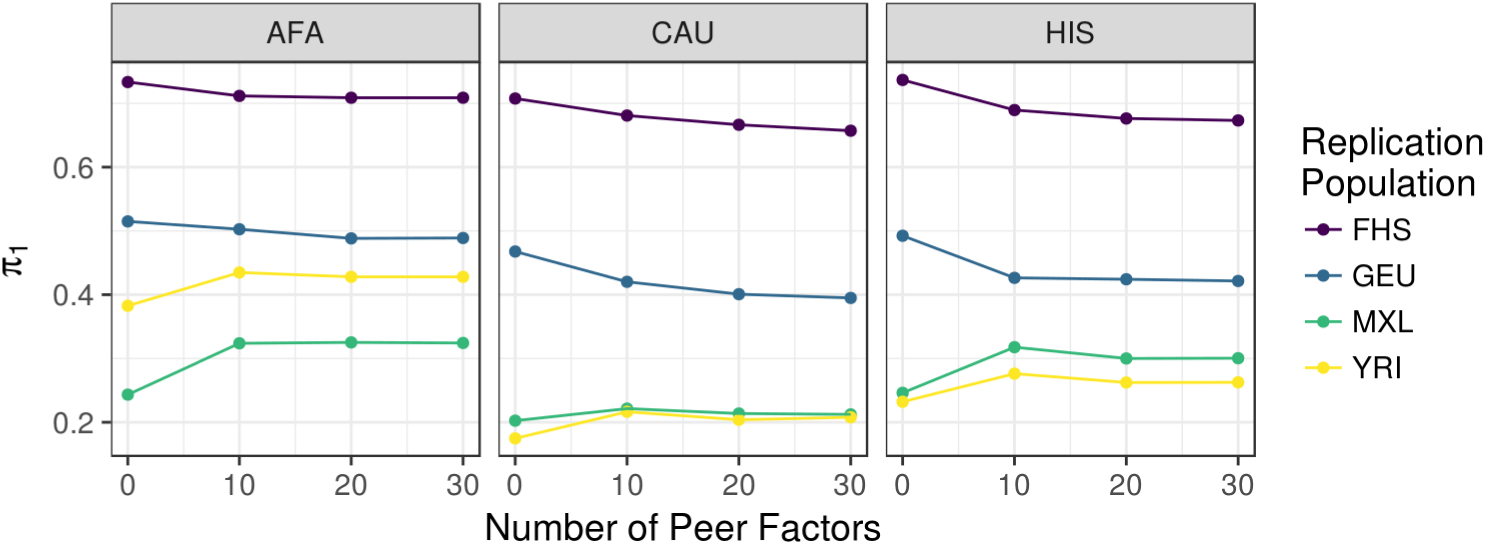
Summary of eQTL analyses in MESA populations. True positive rate π_1_ statistics [29] for cis-eQTLs are plotted vs. the number of PEER factors used to adjust for hidden confounders in the expression data of both discovery and replication populations. The MESA discovery population is listed in the gray title box and the color of the each line represents each replication population. Higher π_1_ values indicate a stronger replication signal. π_1_ is calculated when the SNP-gene pair from the discovery population is present in the replication population. All models shown included 3 genotypic principal components. AFA = MESA African American, CAU = MESA European American, HIS = MESA Hispanic American, FHS = Framingham Heart Study, GEU = Geuvadis, MXL = Mexicans in Los Angeles, YRI = Yoruba in Ibadan, Nigeria.

As expected, the sample size of the discovery population influences the number of eQTLs mapped (Table 1). Hundreds of thousands to millions of SNPs were found to associate with gene expression (eSNPs) and most genes had at least one associated variant (eGenes) at FDR *<* 0.05, with the absolute numbers correlating with sample size (Table 1). Cis-eQTL summary statistics for each population are available at https://github.com/WheelerLab/DivPop.

**Table 1.** cis-eQTL (FDR *<* 0.05) counts across MESA populations

| Population | number eSNPs | number eGenes |
| --- | --- | --- |
| AFA (n=233) | 412,450 | 6837 |
| HIS (n=352) | 890,100 | 7974 |
| CAU (n=578) | 1,290,814 | 7925 |
| AFHI (n=585) | 1,126,620 | 8628 |
| ALL (n=1163) | 1,652,365 | 8877 |

Linear additive models were adjusted for 3 genotypic principal components and 10 PEER factors. FDR = Benjamini-Hochberg false discovery rate. AFA = African American, HIS = Hispanic American, CAU = European American, AFHI = AFA and HIS, ALL = AFA, HIS, and CAU.

### Genetic effect size correlations between populations reflect shared ancestry proportions

We estimated the local (cis-region SNPs) heritability (h^2^) for each gene and the genetic correlation (rG) between genes in each MESA population. We used the average information-REML algorithm implemented in GCTA [30, 31] to estimate rG, which is constrained between −1 and 1 for each gene (See Methods). As in Brown et al. [32], the sample sizes for gene expression data are too small for obtaining accurate point estimates of rG for each gene. However, the large number of genes allow us to obtain accurate estimation of the global mean rG between populations. The population pair with the highest mean rG was CAU and HIS, followed by AFA and HIS, and the least correlated pair was AFA and CAU (Table 2). Genes with larger h^2^ estimates in at least one population tended to have larger rG estimates with lower standard errors (Fig. 2A, S4 Fig). As the h^2^ threshold for inclusion increases, the mean rG between populations also increases (Fig. 2B). The same pattern is observed when the h^2^ estimates are normalized by the number of SNPs in the gene (S4 Fig).

**Table 2.** Genetic correlation (rG) between MESA populations

| pop pair | mean rG | SE rG | genes that converged |
| --- | --- | --- | --- |
| AFA-CAU | 0.46 | 0.0080 | 9209 |
| AFA-HIS | 0.57 | 0.0076 | 9313 |
| HIS-CAU | 0.62 | 0.0072 | 9490 |

rG was estimated using a bivariate restricted maximum likelihood (REML) model implemented in GCTA.

To verify that our rG analysis did not contain any small sample-size biases, we simulated gene expression phenotypes in each population with the same local h^2^ distributions as the real data. For ten sets of simulated gene expression phenotypes, we estimated rG between populations and compared the simulated results to the observed results. While the mean rG ranged from 0.46-0.62 in the observed data, the mean rG in the simulated data was near zero with similar numbers of genes at −1 and 1(Fig. 2C).

**Table 3.** Proportion of genes with greater lasso (*α* = 1) model predictive performance (R^*2*^) compared to elastic net models with different mixing parameter (*α*) values.

| Population | elastic net ($\alpha = 0.05$) | elastic net ($\alpha = 0.5$) |
| --- | --- | --- |
| AFA | 1497/2517 (0.595) | 1342/2567 (0.523) |
| CAU | 2213/3858 (0.574) | 1943/3867 (0.502) |
| HIS | 1950/3529 (0.553) | 1763/3529 (0.500) |

**Fig 2.**
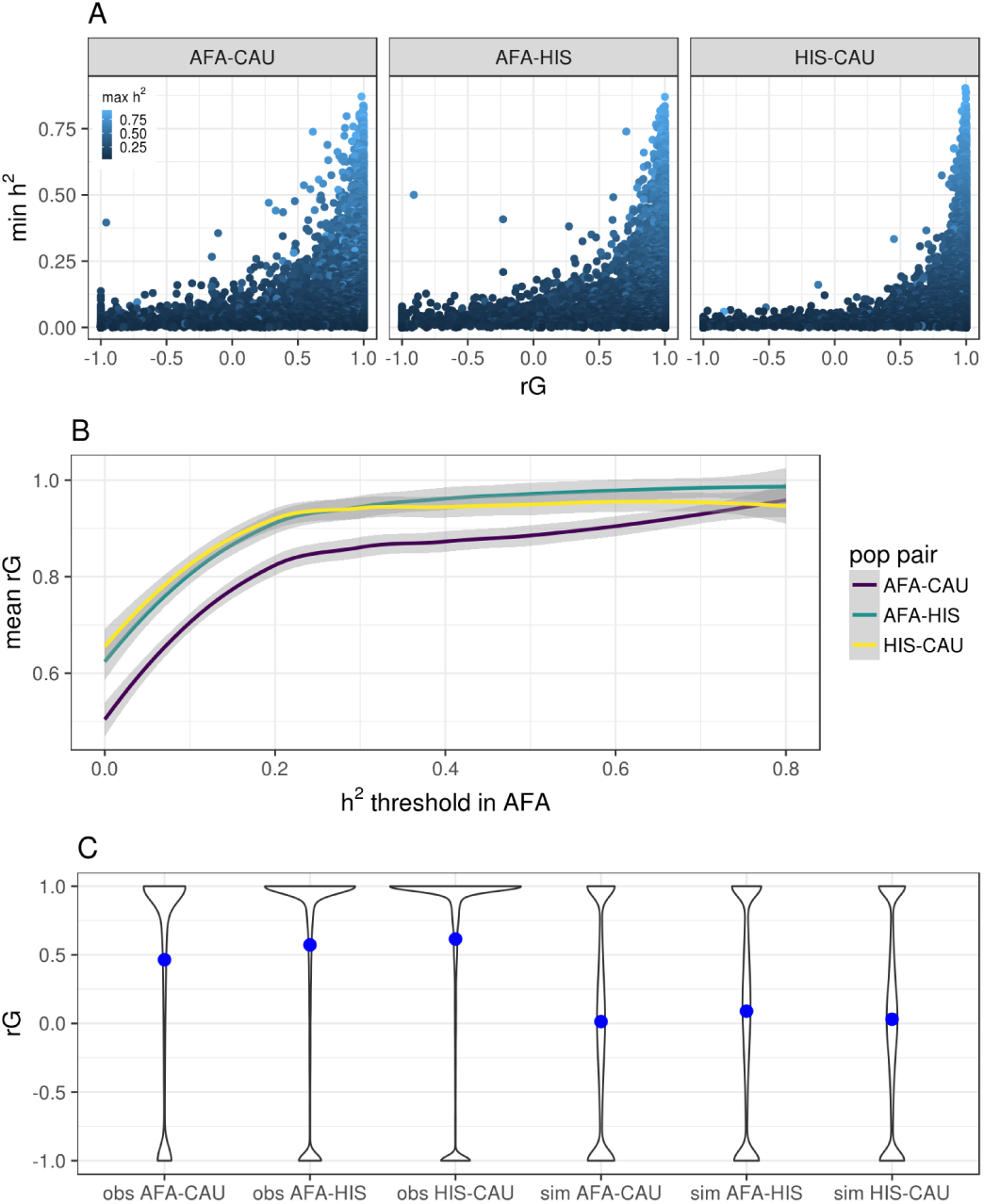
Genetic correlation (rG) of gene expression between MESA populations. (**A**) Pairwise population comparison of heritability (h^2^) and rG for each gene. The y-axis is the minimum h^2^, the x-axis is the genetic correlation, and the points are colored according to the maximum h^2^ between the populations titling each plot. (**B**) Comparison of the genetic correlation between pairwise MESA populations and the subset of genes with h^2^ greater than a given threshold in the AFA population. (**C**) Violin plots of the observed results (obs) compared to simulated expression data (sim) with the same h^2^ distributions. The blue points represent the mean rG across genes for the population pair. The most correlated populations are CAU and HIS and the least correlated populations are AFA and CAU. Note more genes have an rG estimate equal to 1in the observed data compared to the simulated data

**Fig 3.**
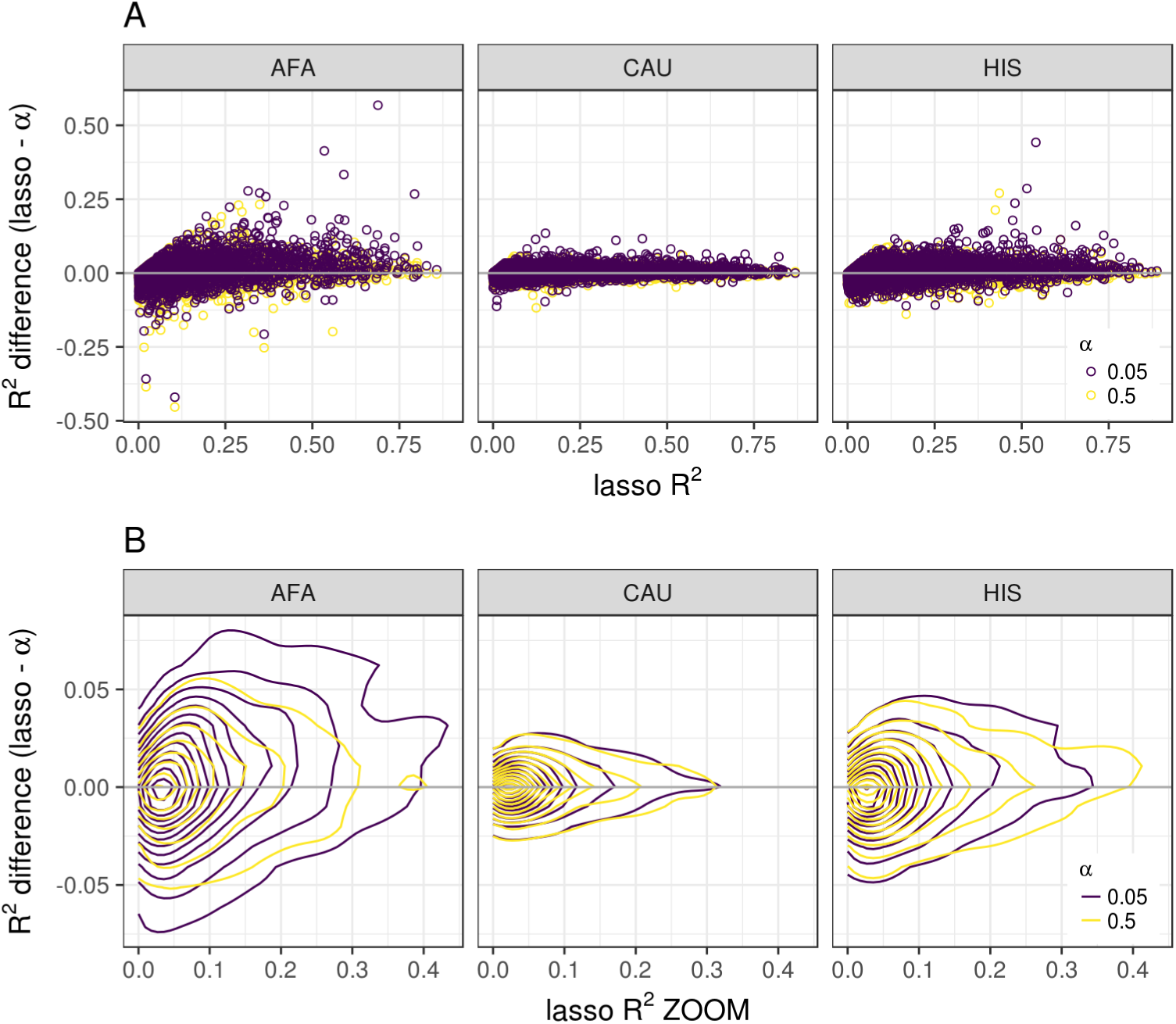
MESA gene expression predictive performance across a range of elastic net mixing parameters. (**A**) The difference between cross-validated R^2^ of lasso and elastic net with mixing parameters 0.05 or 0.5 is compared to the lasso R^2^ across genes in MESA populations AFA, HIS, and CAU. (**B**) Zoomed in plot of **A** using contour lines from two-dimensional kernel density estimation to visualize where the points are concentrated. The R^2^ difference values (y-axis) with a mixing parameter α = 0.5 are closer to zero indicating that they perform similarly to the lasso model. The values with a mixing parameter α = 0.05 are above zero indicating that they perform worse than the lasso model.

### Models with a sparse component outperform polygenic models for gene expression

We examined the prediction performance of a range of models using elastic net regularization [33] to characterize the genetic architecture of gene expression in each population. The mixing parameter (*α*) of elastic net ranges from 0-1. Models with *α* near 0 assume a more polygenic architecture and models with *α* near 1assume a more sparse architecture. We used nested cross-validation to compute the coefficient of determination R^2^ as our measure of model performance across three mixing parameters (*α* = 0.05, 0.5, 1). The model with *α* = 1is equivalent to least absolute shrinkage and selection operator (lasso) regression [34]. When we compared the R^2^ values for each gene between models, more genes had a higher R^2^ with the lasso model (*α* = 1) than the most polygenic model tested (*α* = 0.05) in each population (Table 3, Fig. 3). The lasso model performed similarly to the mixture model (*α* = 0.5), indicating elastic net is somewhat robust to *α* choice as long as a sparse component is included (Table 3, Fig. 3).

In addition to elastic net, we used Bayesian Sparse Linear Mixed Modeling (BSLMM) [35] to estimate if the local genetic contribution to gene expression is more polygenic or sparse. This approach models the genetic contribution of the trait as the sum of a sparse component and a polygenic component. BSLMM estimates the total percent variance explained (PVE) and the parameter PGE, which represents the proportion of the genetic variance explained by sparse effects. We found that for highly heritable genes (high PVE), the sparse component (PGE) is large; however, for genes with low PVE, we are unable to determine whether the sparse or polygenic component is predominant (S5 Fig). We also estimated heritability (h^2^) using a linear mixed model (LMM) [30] and Bayesian variable selection regression (BVSR) [36], which assume a polygenic and sparse architecture, respectively. It has previously been shown that BVSR performs similarly to BSLMM when the simulated architecture is sparse, but BVSR performs poorly compared to BSLMM when the simulated architecture includes a polygenic component [35]. BSLMM outperforms both LMM and BVSR in each population (S5 Fig). However, BSLMM and BVSR show greater correlation, providing further support that the sparse component dominates in the MESA cohorts (S5 Fig).

### Differences in predictive performance are due to allele frequency differences between populations

We then compared each population’s gene expression predictive performance as measured by cross-validated coefficient of determination (R^2^). We first fit elastic net models (*α* = 0.5) using 3 genotypic PCs and gene expression levels adjusted by 0, 10, 20 or 30 PEER factors in each population. Predictive performance was higher when we used 10 PEER factors compared to no PEER factor adjustment (S6 Fig). Seeing little difference between models with 10 or more PEER factors within populations (S6 Fig), we compared predictive performance between populations using the elastic net models with 10 PEER factors. The Spearman correlation (*ρ*) between CAU and HIS model performance is highest (*ρ* = 0.778), followed by AFA and HIS (*ρ* = 0.663). The lowest correlation between two populations was AFA and CAU with *ρ* = 0.586 (Fig. 4A). These correlation relationships mirror the European and African admixture proportions in the MESA HIS and AFA cohorts (S1 Fig).

Because the sample sizes between MESA populations differed (Table 1), we randomly selected 233 individuals from CAU and HIS and fit elastic net models with these downsampled populations to match the AFA sample size. Predictive performance R^2^ is highly correlated between the full and downsampled populations (*ρ >* 0.96). A handful of genes that are better predicted with the full sample size (S7 Fig). Also, the between population correlations showed the same trend when all populations had the same sample size, with CAU and HIS the most correlated, followed by AFA and HIS (S7 Fig).

There are many genes that are well predicted in both populations and poorly predicted in both populations. We found the best predicted gene, *TACSTD2*, was the same across each population with an R^2^ > 0.86 in each population. On the other hand, there are some genes that are well predicted in one population, but poorly predicted in the other and vice versa (Fig. 4A).

To test the hypothesis that allele frequency differences between populations are influencing predictive power, we performed a fixation index (F_ST_) analysis. For each population pair, we calculated the the mean F_ST_ for SNPs in each gene expression prediction model. Gene models with an absolute value R^2^ difference between populations greater than 0.05 had significantly higher mean F_ST_ distribution than those with a smaller difference (Wilcoxon *P* = 2.7×10^−66^). The significant increase in mean F_ST_ was robust across R^2^ difference thresholds (Fig. 4C). Similar significant differences were observed when the SNP F_ST_ values were weighted by elastic net model betas across R^2^ difference thresholds from 0.05-0.3 (S8 Fig).

### Gene expression prediction improves when training set has similar ancestry to test set

In order to further compare gene expression prediction model performance between populations, using the models built in each MESA population, we predicted gene expression in our replication populations: FHS, GEU, MXL, and YRI. We calculated the true positive rates (π_1_statistics) [29] for predicted vs. observed expression in each replication cohort when different numbers of gene models were included based on MESA predictive performance (Table 4, Fig. 5). As expected, true positive rates were higher across model training populations for the largest replication population, FHS. For GEU, which includes European and African ancestry individuals, the best performing models were trained using all the MESA individuals (ALL, Fig. 5). Prediction in YRI was best using AFA or AFHI models and prediction in MXL was optimal using the CAU models (Fig. 5). These results demonstrate that when comparing predicted expression levels to the observed, a balance of the training population with ancestry most similar to the test population and total sample size leads to optimal predicted gene expression.

**Fig 4.**
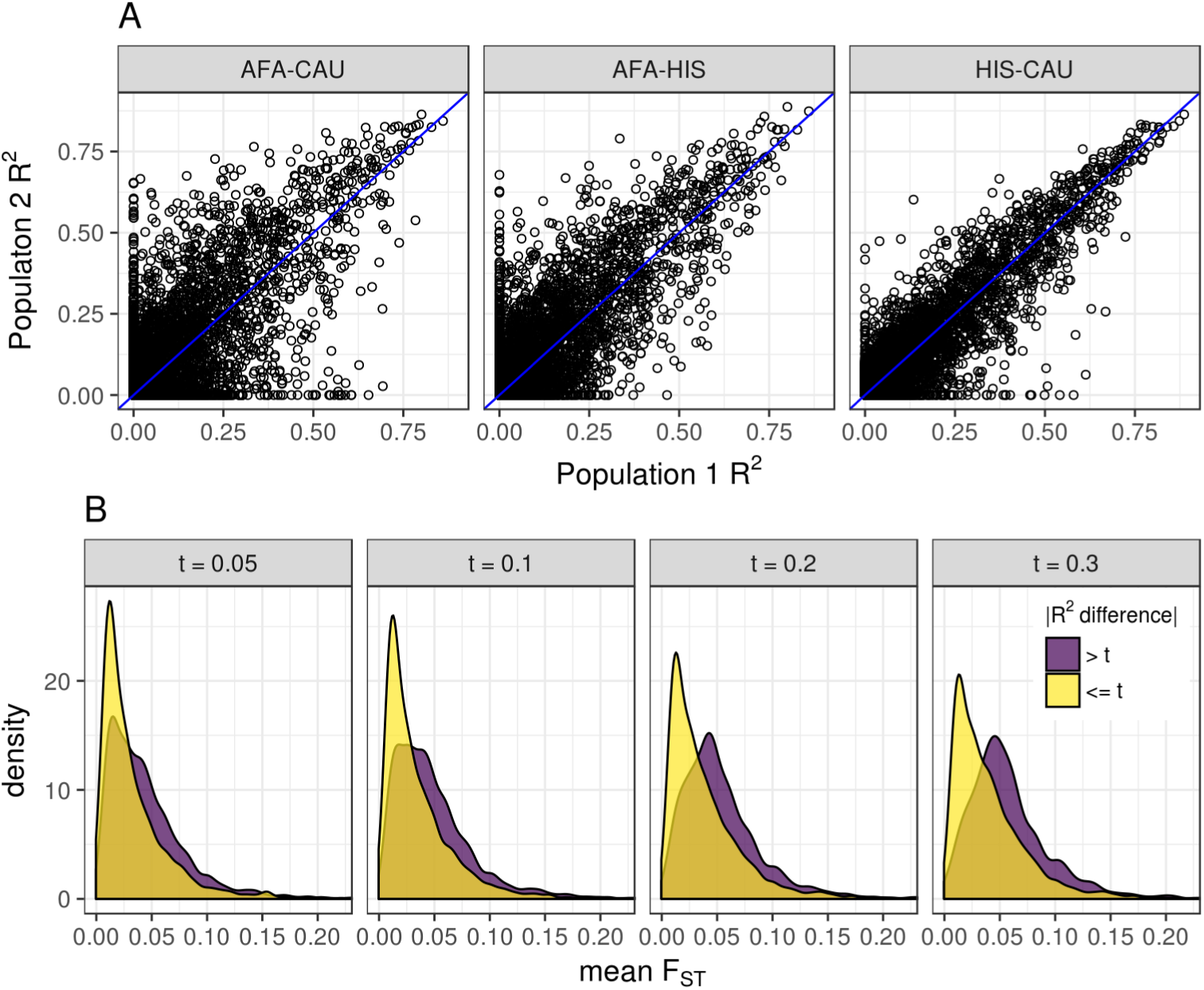
Allele frequency differences lead to gene expression predictive performance differences between populations. (**A**) Comparison of predictive performance for each gene (R^2^) between each pair of populations. Predictive performance (R^2^) was measured within each population using nested cross-validation. In each gray title box, population 1is listed first and population 2 is listed second. The identity line is shown in blue. The pairwise Spearman correlations (*ρ*) between genes are AFA-CAU: *ρ* = 0.762, AFA-HIS: *ρ* = 0.84, HIS-CAU: *ρ* = 0.92. (**B**) Comparison of mean FST between gene models with large (> t) and small (*<*= t) differences in predictive performance R^2^. Mean FST of SNPs in each gene expression prediction model between all pairwise populations was calculated. The gene groups with the larger absolute value R^2^ difference between populations had significantly larger mean FST at each difference threshold, t (Wilcoxon rank sum tests, *P* < 2.2 *×;* 10^−16^).

**Fig 5.**
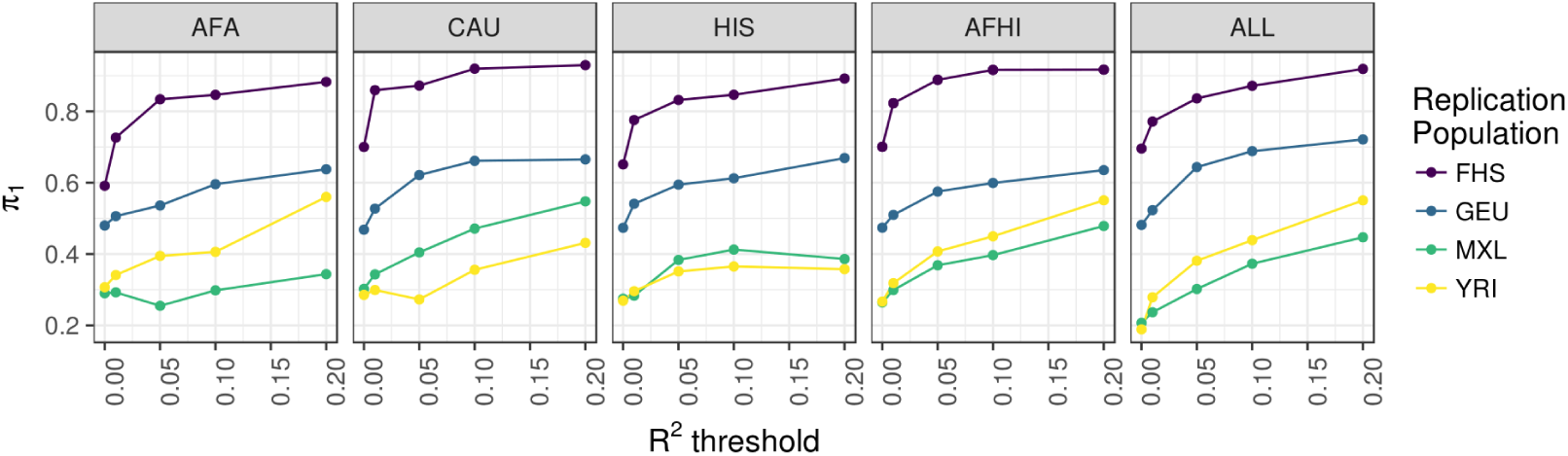
Predictive performance in independent test cohorts across MESA population models.

True positive rate π_1_statistics [29] for replication cohort prediction are plotted vs. the training population model predictive performance R^2^. The MESA training population is listed in the gray title box and the color of each line represents each replication population. Higher π_1_values indicate a higher true positive rate of predicted expression in the replication cohort using the MESA model vs. observed expression in the replication cohort. All models shown included 3 genotypic principal components and 10 PEER factors. AFA = MESA African American, CAU = MESA European American, HIS = MESA Hispanic American, FHS = Framingham Heart Study, GEU = Geuvadis, MXL = Mexicans in Los Angeles, YRI = Yoruba in Ibadan, Nigeria.

**Table 4.** Number of genes with models at different R^2^ thresholds.

| Model | $R^2 \geq 0$ | $R^2 \geq 0.01$ | $R^2 \geq 0.05$ | $R^2 \geq 0.1$ | $R^2 \geq 0.2$ |
| --- | --- | --- | --- | --- | --- |
| AFA | 3486 | 3006 | 2153 | 1584 | 910 |
| HIS | 4457 | 3879 | 2704 | 1913 | 1152 |
| CAU | 4901 | 4128 | 2753 | 1921 | 1149 |
| AFHI | 5778 | 4926 | 3303 | 2304 | 1308 |
| ALL | 6896 | 5672 | 3532 | 2407 | 1336 |

The number of genes in MESA (AFA = African American, HIS = Hispanic American, CAU = European American, AFHI = AFA and HIS, ALL = AFA, HIS, and CAU) as training sets to predict gene expression

### Gene-based association using multiethnic predictors

Gene-based association methods like PrediXcan, TWAS, and S-PrediXcan have been developed to use genotype data to discover genes whose predicted expression is associated a phenotype of interest [20, 21, 37]. To date, most predicted expression models available for these methods were trained in European ancestry cohorts. We used the five MESA models with S-PrediXcan [37] and publicly available multiancestry GWAS summary statistics from a large asthma study by the Trans-National Asthma Genetic Consortium (TAGC) [38]. While all MESA models performed similarly, the top genes differed across models (Fig. 6, S1 Table). Many genes identified by S-PrediXcan were not previously implicated in TAGC GWAS [38] (Table 5, S1 Table). Two of the genes that associated with asthma using the ALL models were not predicted in CAU and thus not even tested, demonstrating the additional information non-European populations may add to studies. They include *C2* (complement C2) and *BLOC1S1*(biogenesis of lysosomal organelles complex 1subunit 1), which are on different chromosomes. Neither gene has been implicated in asthma GWAS before, but both are associated with age-related macular degeneration, another inflammation-related disease [39]. All summary statistics from our S-PrediXcan analyses are in S2 Table.

**Fig 6.**
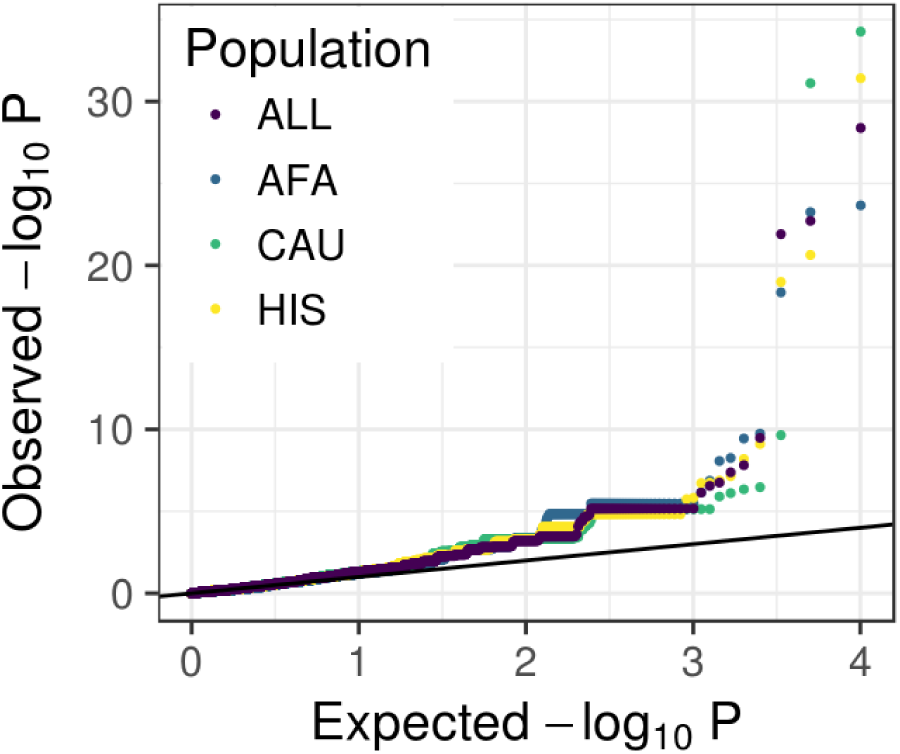
Comparison of S-PrediXcan results using gene expression prediction models from different MESA populations and summary statistics from a multiancestry GWAS of asthma.

Summary statistics were retrieved from the GWAS Catalog for the Trans-National Asthma Genetic Consortium study [38]. Q-Q plots of S-PrediXcan results using models built in each population.

**Table 5.** Summary of S-PrediXcan results using MESA models in a multiancestry GWAS of the asthma [38].

| Model | Bonferroni threshold | significant genes | also significant in GWAS | also significant using CAU |
| --- | --- | --- | --- | --- |
| AFA | 1.5e-5 | 10 | 4 | 6/10 |
| HIS | 1.1e-5 | 14 | 4 | 4/12 |
| CAU | 1.1e-5 | 17 | 7 | NA |
| AFHI | 9.2e-6 | 16 | 4 | 8/14 |
| ALL | 8.1e-6 | 17 | 5 | 10/15 |

## Discussion

We compared three MESA populations (AFA, HIS, and CAU) to better understand the genetic architecture of gene expression in diverse populations. We optimized predictors of gene expression using elastic net regularization and found that models with a sparse component outperform polygenic models. Between populations, the genetic correlation of gene expression is higher when continental ancestry proportions are more similar. We identified genes that are better predicted in one population and poorly predicted in another due to allele frequency differences. We tested our predictors developed in MESA in independent cohorts and found that the best prediction of gene expression occurred when the training set included individuals with similar ancestry to the test set.

As seen in other studies [18, 21, 40], we show models with a sparse component outperform polygenic only models for gene expression prediction across populations. Thus, the genetic architecture of gene expression for many genes has a substantial sparse component. Notably, some genes do perform better in more polygenic models as shown here (genes below the horizontal zero line in Fig. 3 and S5 Fig) and in Zeng et al. [41]. Larger sample sizes may reveal an additional polygenic component that may improve prediction for some genes. However, the population with the largest sample size (CAU) showed the least variability between models (Fig. 3), suggesting that a more polygenic model does not add much to the predictive performance of a sparse model with fewer predictors. Thus, to balance these observations, we recommend using models that include a mixture of polygenic and sparse components like elastic net (α ≃ 0.5) [33], BSLMM [35], and latent Dirichlet process regression [41].

We estimated the genetic correlation between each population pair for each gene. Populations with more shared ancestry as defined by clustering of genotypic principal components showed higher mean correlation across genes (Fig. S1 Fig, Table 2). As estimated heritability of genes increase, the mean genetic correlation between populations also increases (Fig. 2B), which indicates the genetic architecture underlying gene expression is similar for the most heritable genes. However, even though prediction across populations is possible for some of the most heritable genes, we define a class of genes where predictive performance drops substantially between populations. We show this drop is due to allele frequency differences (larger F_ST_) between populations. We tested our predictive gene expression models built in the MESA populations in several replication populations. As expected, the YRI gene expression prediction was best when using the AFA, AFHI, or ALL training sets, which each include individuals with African-ancestry admixture (Fig. 5). The best gene expression prediction for MXL was with the CAU training set, which may reflect the lack of recent African ancestry in MXL [6] compared to the MESA HIS population (S1 Fig). For GEU, the best MESA prediction population was ALL, which indicates that multi-ethnic cohorts like GEU benefit from a pooled training set containing individuals of diverse ancestries. Thus, it may be beneficial to build gene expression models using training populations with a similar allele frequency spectrum to that of the test cohort taking into account SNPs that are interrogated in both populations. A similar cohort-specific strategy was used to increase power to detect genes associated with warfarin dose using PrediXcan in African Americans [42].

We applied S-PrediXcan using our MESA models to summary statistics from a multiancestry GWAS of asthma [38]. We found several novel and previously reported genes significantly associated with asthma (Table 5, S1 Table). Of the genes not implicated in the Demenais et al. GWAS [38], most were associated with inflammation-related diseases in the GWAS Catalog [1]. We found increased predicted *ADORA1* expression significantly associated with increased asthma risk in 4/5 MESA models tested (S2 Table). While *ADORA1* was not significant in Demenais et. al. [38], the gene has previously been reported to associate with asthma in a study investigating the relationships between phenotypes, which also found that immune-related disease associations cluster together [43]. Similar inflammation mechanisms could explain why two genes (*C2* and *BLOC1S1*) previously associated with age-related macular degeneration [39] might also be implicated in asthma as shown here.

Predictive models of gene expression developed in this study and performance statistics are made publicly available at https://github.com/WheelerLab/DivPop for use in future studies of complex trait genetics across diverse populations. As in our S-PrediXcan analysis of asthma, multiancestry transcriptome integration may reveal new genes not implicated in European only studies. Inclusion of diverse populations in complex trait genetics is crucial for equitable implementation of precision medicine.

## Materials and methods

This work was approved by the Loyola University Chicago Institutional Review Board.

### Genomic and transcriptomic training data

#### The Multi-Ethnic Study of Atherosclerosis (MESA)

MESA includes 6814 individuals consisting of 53% females and 47% males between the ages of 45-84 [24]. The individuals were recruited from 6 sites across the US (Baltimore, MD; Chicago, IL; Forsyth County, NC; Los Angeles County, CA; northern Manhattan, NY; St. Paul, MN). MESA cohort population demographics were 39% European American (CAU), 22% Hispanic American (HIS), 28% African American (AFA), and 12% Chinese American (CHN). Of those individuals, RNA was collected from CD14+ monocytes from 1264 individuals across three populations (AFA, HIS, CAU) and quantified on the Illumina Ref-8 BeadChip [25]. Individuals with both genotype (dbGaP: phs000209.v13.p3) and expression data (GEO: GSE56045) included 234 AFA, 386 HIS, and 582 CAU. Illumina IDs were converted to Ensembl IDs using the RefSeq IDs from MESA and gencode.v18 (gtf and metadata files) to match Illumina IDs to Ensembl IDs. If there were multiple Illumina IDs corresponding to an Ensembl ID, the average of those values was used as the expression level.

### Genomic and transcriptomic test data

#### Stranger et al. HapMap data

We obtained lymphoblastoid cell line (LCL) microarray transcriptome data from Stranger et al. [14] for HapMap populations of interest, including 45 Mexican ancestry individuals in Los Angeles, CA, USA (MXL) and 107 Yoruba individuals in Ibadan, Nigeria (YRI) (Illumina Sentrix Human-6 Expression BeadChip version 2, Array Express: E-MTAB-264). We obtained genotype data from the 1000 Genomes Project (phase3 v5a 20130502) [44]. HapMap genotypes in individuals not sequenced through the 1000 Genomes Project were imputed using the Michigan Imputation Server for a total of 6-13 million SNPs per population, after undergoing quality control [45]. These imputed samples were then merged with the individuals that were previously sequenced, filtering the SNPs (imputation R^2^ > 0.8, MAF > 0.01, HWE p > 1e-06).

#### Geuvadis Consortium (GEU)

We obtained RNA sequencing transcriptome data from the Geuvadis Consortium (GEU) at https://www.ebi.ac.uk/arrayexpress/experiments/E-GEUV-1/ and genotype data from the 1000 Genomes Project (phase3 v5a 20130502) [26, 44]. The GEU cohort includes 78 Utah residents with Northern and Western European ancestry, 89 Finnish from Finland, 85 British from England and Scotland, 92 Toscani in Italy and 77 Yoruba in Ibadan, Nigeria individuals [26].

#### Framingham Heart Study (FHS)

We obtained genotype and exon expression array (Affymetrix Human Exon 1.0 ST microarray) data [27] through application to dbGaP accession phs000007.v29.p1. Genotype imputation and gene level quantification were performed by our group previously [18], leaving 4838 European ancestry individuals with both genotypes and observed gene expression levels for analysis.

### Quality control of genomic and transcriptomic data

We imputed genotypes in the MESA populations using the Michigan Imputation Server and 1000 genomes phase 3 v5 reference panel and Eagle v2.3. Reference populations were EUR for CAU and mixed population for AFA and HIS [44–46]. The results were filtered by R^2^ *<* 0.8, MAF > 0.01, and ambiguous strand SNPs were removed. This left 9,352,383 SNPs in AFA, 7,201,805 SNPs in HIS, and 5,559,636 SNPs in CAU for further analysis. Quality control and cleaning of the genotype data was done using PLINK (https://www.cog-genomics.org/plink2). SNPs were filtered by call rates less than 99%. Prior to IBD and principal component (PC) analysis, SNPs were LD pruned by removing 1SNP in a 50 SNP window if r^2^ > 0.3. One of a pair of related individuals (IBD > 0.05) were removed. Pruned genotypes were merged with HapMap populations and EIGENSTRAT [47] was used to perform PC analysis both across (Fig. S1 Fig) and within populations. Final sample sizes for each population post quality control are AFA = 233, HIS = 352, and CAU = 578. We used 5-7 million non-LD pruned SNPs per population post quality control. PEER factor analysis within each population was performed on the expression data using the peer R package in order to correct for potential batch effects and experimental confounders [48].

### eQTL analysis

We used Matrix eQTL [49] to perform a genome-wide cis-eQTL analysis in each population separately (AFA, HIS, CAU), in the AFA and HIS combined (AFHI), and in all three populations combined (ALL). We used SNPs with MAF > 0.01and defined cis-acting as SNPs within 1Mb of the transcription start site (TSS). We tested a range of linear regression models with 0, 3, 5, or 10 within population genotypic PC covariates and 0, 10, 20, or 30 within population PEER factors [28]. The false discovery rate (FDR) for each SNP was calculated using the Benjamini-Hochberg procedure. We estimate the pairwise population eQTL true positive rates with π_1_statistics using the qvalue method [17, 29]. π_1_is the expected true positive rate and was estimated by selecting the SNP-gene pairs with FDR *<* 0.05 in each discovery cohort (MESA) and examining their P value distribution in each replication cohort (FHS, GEU, MXL, YRI). *π*_0_ is the proportion of false positives estimated by assuming a uniform distribution of null P values and π_1_= 1 *− π*_0_ [29].

### Genetic correlation analysis

We performed eQTL effect size comparisons between populations using Genome-wide Complex Trait Analysis (GCTA) software [30]. We performed a bivariate restricted maximum likelihood (REML) analysis to estimate the genetic correlation (rG) between each pair of MESA populations for each gene [31]. As in the eQTL analysis, we compared cis-region (within 1Mb) SNPs for each gene. In our implementation, the models can be written as

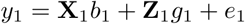

for population 1and

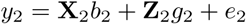

where *y*_1_and *y*_2_ are vectors of gene expression values, *b*_1_and *b*_2_ are vectors of fixed effects, *g*_1_and *g*_2_ are vectors of random polygenic effects, and *e*_1_and *e*_2_ are residuals for populations 1and 2, respectively. **X** and **Z** are incidence matrices for the effects *b* and *g*, respectively. The variance covariance matrix (**V**) is defined as

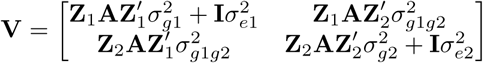

where **A** is the genetic relationship matrix based on SNP information [30], **I** is an identity matrix, *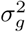* is the genetic variance, 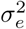 the is residual variance, and *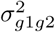* is the covariance between *g*_1_and *g*_2_. In our models, the residual covariance component is ignored because no individual belongs to two populations. We used the average information-REML algorithm implemented in GCTA [31] to estimate rG, which is constrained between −1 and 1for each gene by bending the variance-covariance matrix to be positive definite.

As in Brown et al. [32], the sample sizes for gene expression data are too small for obtaining accurate point estimates of rG for each gene. However, the large number of genes allow us to obtain accurate estimation of the global mean rG between populations. To verify that our rG analysis did not contain any small sample-size biases, we simulated gene expression phenotypes in each population with the same local heritability (*h*^2^) distributions as the real data. Effect sizes of cis-region SNPs for each gene were randomly generated from a standard normal distribution such that the individual population *h*^2^ estimate would be the same as the observed data. For ten sets of simulated gene expression phenotypes, we estimated rG between populations and compared the simulated results to the observed results (Fig. 2).

### Prediction model optimization

We used the glmnet R package [33] to fit an elastic net model to predict gene expression from cis-region SNP genotypes. The elastic net regularization penalty is controlled by the mixing parameter alpha, which can vary between ridge regression (*α* = 0) and lasso (*α* = 1, default). A gene with the optimal predictive performance when *α* = 0 has a polygenic architecture, whereas a gene with optimal performance when *α* = 1has a sparse genetic architecture. In the MESA cohort we tested three values of the alpha mixing parameter (0.05, 0.5, and 1) and a range of PEER factors (0, 10, 20, 30) for optimal prediction of gene expression of 10,143 genes for each population alone (AFA, CAU, HIS), AFA and HIS combined (AFHI), and all three populations combined (ALL). We used the PredictDB pipeline developed by the Im lab to preprocess, train, and compile elastic net results into database files to use as weights for gene expression prediction [37]. We quantified the predictive performance of each model via nested cross-validation. We split the data into 5 disjoint folds, roughly equal in size, and for each fold, we calculated a 10-fold cross-validated elastic net model in 4/5 of the data where the lambda tuning parameter is cross-validated. Then, using predicted and observed gene expression, we calculate the coefficient of determination (R^2^) for how the model predicts on the held-out fold. We report the mean R^2^ over all 5 folds as our measure of model performance. R^2^ is defined as

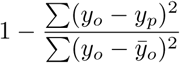

where *y*_*o*_ is observed expression, *y*_*p*_ is predicted expression, and 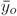 is the mean of observed expression. See https://github.com/WheelerLab/DivPop.

We used the software GEMMA [50] to implement Bayesian Sparse Linear Mixed Modeling (BSLMM) [35] for each gene with 100K sampling steps per gene. BSLMM estimates the PVE (the proportion of variance in phenotype explained by the additive genetic model, analogous to the h^2^ estimated in GCTA) and PGE (the proportion of genetic variance explained by the sparse effects terms where 0 means that genetic effect is purely polygenic and 1means that the effect is purely sparse). From the second half of the sampling iterations for each gene, we report the median and the 95% credible sets of the PVE and PGE. We also estimated heritability (h^2^) using a linear mixed model (LMM) implemented in GCTA [30] and Bayesian variable selection regression (BVSR) [36], which assume a polygenic and sparse architecture, respectively. We used the software piMASS for Bayesian variable selection regression (BVSR) [36]. For each gene, we used 10,000 burn-in steps and 100,000 sampling steps in the BVSR Markov chain Monte Carlo algorithm. From the output of every 10 sampling steps, we report the median re-estimated PVE based on sampling posterior effect sizes.

### Comparing prediction models between MESA populations

We calculated the fixation index (F_ST_) [51] for each SNP between each pair of populations using PLINK. Then, for each gene expression prediction model, we calculated both the mean F_ST_ and weighted average F_ST_ for SNPs in the model. In the weighted average calculation, F_ST_ values were multiplied by the elastic net model beta value to give SNPs with larger effect sizes more weight. We compared mean and weighted average F_ST_ values between genes with divergent predictive performance and genes with similar predictive performance between populations using Wilcoxon rank sum tests. To test for robustness across thresholds, we varied the absolute value R^2^ difference threshold to define the divergent and similar groups from 0.05-0.3.

### Testing prediction models in independent replication cohorts

Using our elastic net models built in MESA AFA, HIS, CAU, AFHI, and ALL (*α* = 0.5 with 10 PEER factors and 3 genotypic PCs), we predicted gene expression from genotypes in independent test populations: FHS, GEU, MXL, and YRI. As for eQTLs, we estimated the pairwise population prediction true positive rates with π_1_statistics using the qvalue method [17, 29]. The Pearson correlation between predicted and observed expression was calculated and the P value distribution of the correlation was evaluated using π_1_statistics. We calculated π_1_values in the test populations using several MESA model predictive performance R^2^ thresholds for gene inclusion (R^2^ = 0, 0.01, 0.05, 0.1, and 0.2).

### S-PrediXcan application of MESA gene expression prediction models

We performed S-PrediXcan [37] with MESA models AFA, HIS, CAU, AFHI, and ALL using publicly available multiancestry GWAS summary statistics from a large asthma study by the Trans-National Asthma Genetic Consortium (TAGC) [38]. TAGC contained 23,948 cases and 118,538 controls with the following ancestry proportions: 127,669 European, 8,204 African, 5,215 Japanese, and 1,398 Latino [38]. The Bonferroni correction threshold used to define significant genes was calculated as P *<* (0.05/gene count).

## Acknowledgments

This work is supported by the NIH National Human Genome Research Institute Academic Research Enhancement Award R15 HG009569 (HEW), start-up funds from Loyola University Chicago (HEW), the Loyola Carbon Undergraduate Research Fellowship (AA), the Loyola Biology Summer Research Fellowship (AB), the Loyola Mulcahy Scholars Program (AB), and R01MH107666 (HKI). MESA and the MESA SHARe project are conducted and supported by the National Heart, Lung, and Blood Institute (NHLBI) in collaboration with MESA investigators. Support for MESA is provided by contracts HHSN268201500003I, N01-HC-95159, N01-HC-95160, N01-HC-95161, N01-HC-95162, N01-HC-95163, N01-HC-95164, N01-HC-95165, N01-HC-95166, N01-HC-95167, N01-HC-95168, N01-HC-95169, UL1-TR-000040, UL1-TR-001079, UL1-TR-001420, UL1-TR-001881, and DK063491. Funding for SHARe genotyping was provided by NHLBI Contract N02-HL-64278. Genotyping was performed at Affymetrix (Santa Clara, California, USA) and the Broad Institute of Harvard and MIT (Boston, Massachusetts, USA) using the Affymetrix Genome-Wide Human SNP Array 6.0. The MESA Epigenomics & Transcriptomics Study was funded by NIA grant 1R01HL101250-01to Wake Forest University Health Sciences (YL). The Framingham Heart Study is conducted and supported by the National Heart, Lung, and Blood Institute (NHLBI) in collaboration with Boston University (Contract No.N01-HC-25195 and HHSN268201500001I). This manuscript was not prepared in collaboration with investigators of the Framingham Heart Study and does not necessarily reflect the opinions or views of the Framingham Heart Study, Boston University, or NHLBI. Funding for SHARe Affymetrix genotyping was provided by NHLBI Contract N02-HL-64278. SHARe Illumina genotyping was provided under an agreement between Illumina and Boston University. Funding for Affymetrix genotyping of the FHS Omni cohorts was provided by Intramural NHLBI funds from Andrew D. Johnson and Christopher J. O’Donnell. Additional funding for SABRe was provided by Division of Intramural Research, NHLBI, and Center for Population Studies, NHLBI. The following datasets were downloaded from dbGaP: phs000363.v12.p9 and phs000342.v13.p9. The funders had no role in study design, data collection and analysis, decision to publish, or preparation of the manuscript.

## Supporting information

**S1 Fig.**
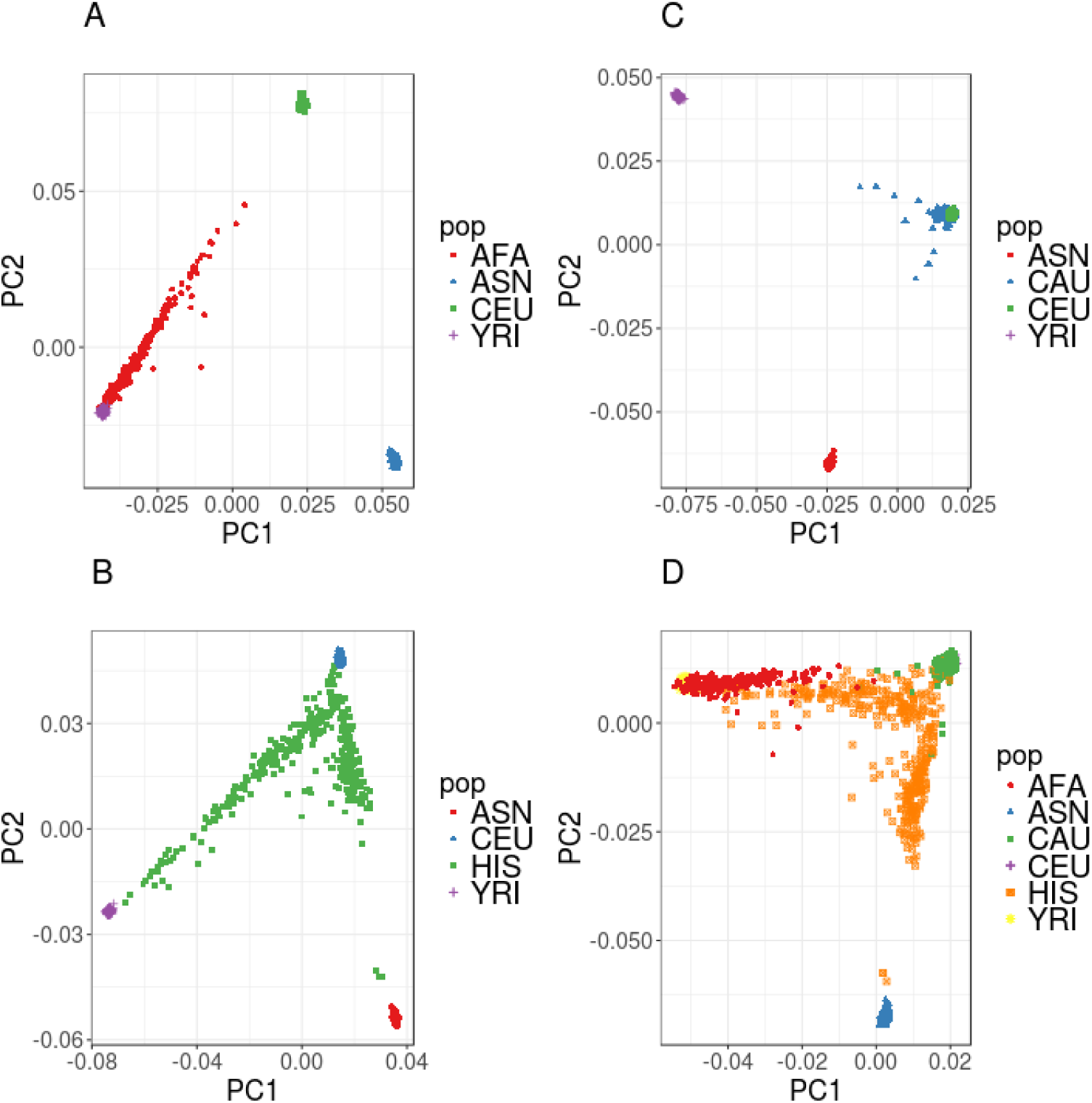
Genotypic principal component (PC) analysis of MESA populations. PC1vs. PC2 plots of each MESA population when analyzed with HapMap populations show varying degrees of admixture. The HapMap populations are defined by the following abbreviations: Yoruba from Ibadan, Nigeria (YRI), European ancestry from Utah (CEU), East Asians from Beijing, China and Tokyo, Japan (ASN). MESA AFA population (red), (**B**) MESA HIS population (green), (**C**) MESA CAU population (blue), (**D**) all MESA populations combined.

**S2 Fig.**
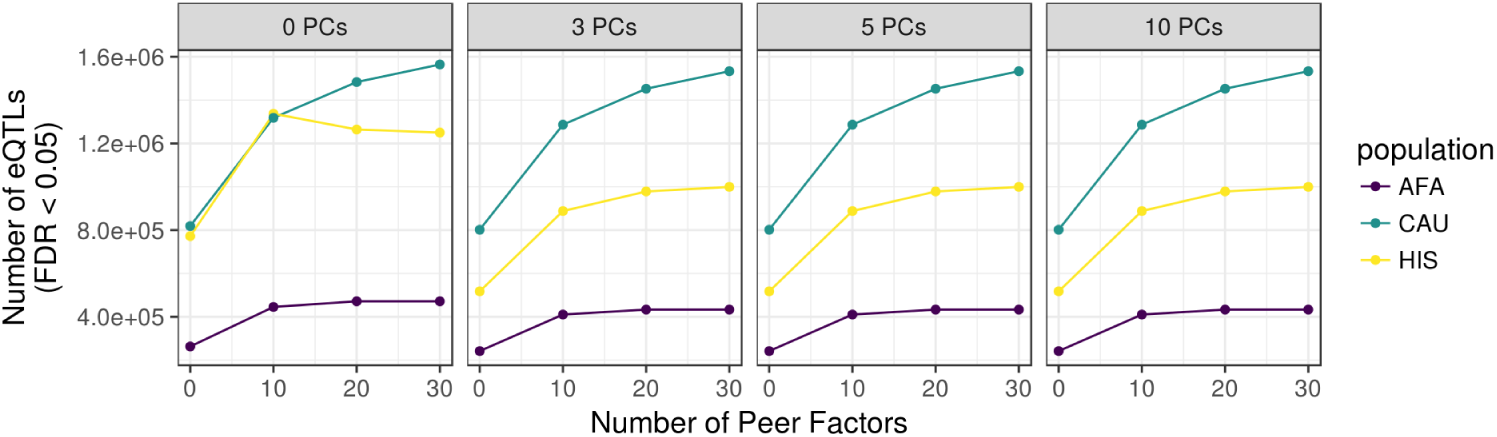
Consistent cis-eQTL results with 3 or more genotypic principal components (PCs). cis-eQTL count (FDR *<* 0.05) vs. the number of PEER factors used to adjust for hidden confounders in the expression data of each MESA population. The number of genotypic PCs is listed in the gray title box and the color of the lines represent each MESA population. Note that all curves with at least 3 genotypic PCs look the same. AFA = MESA African American, CAU = MESA European American, HIS = MESA Hispanic American.

**S3 Fig.**
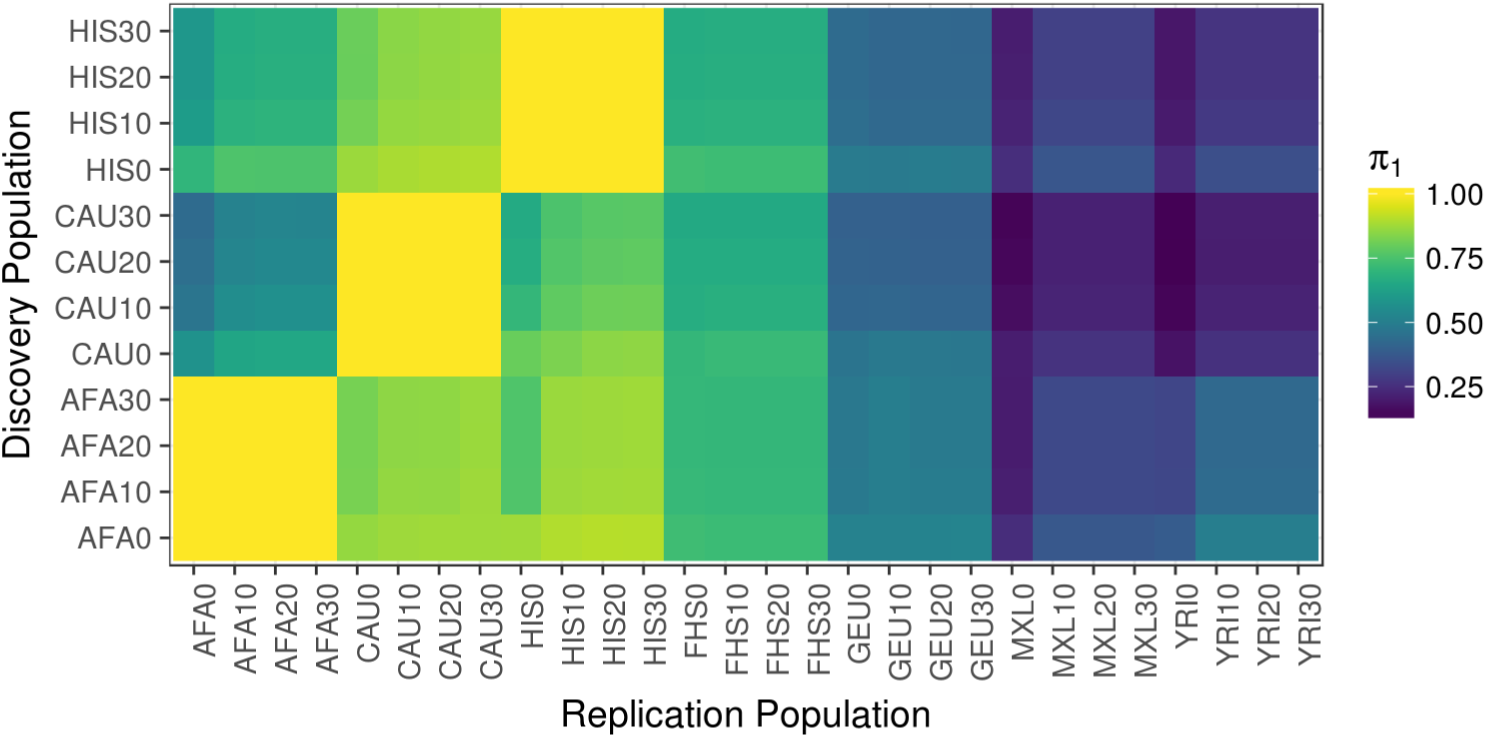
Pairwise cis-eQTL true positive rates across all PEER factor combinantions. True positive rate π_1_statistics [29] for cis-eQTLs are plotted comparing each Discovery Population to each Replication Population. The number after each population abbreviation is the number of PEER factors used to adjust for hidden confounders in the expression data. Higher π_1_values indicate a stronger replication signal. π_1_is calculated when the SNP-gene pair from the discovery population is present in the replication population. All models shown included 3 genotypic principal components. AFA = MESA African American, CAU = MESA European American, HIS = MESA Hispanic American, FHS = Framingham Heart Study, GEU = Geuvadis, MXL = Mexicans in Los Angeles, YRI = Yoruba in Ibadan, Nigeria.

**S4 Fig.**
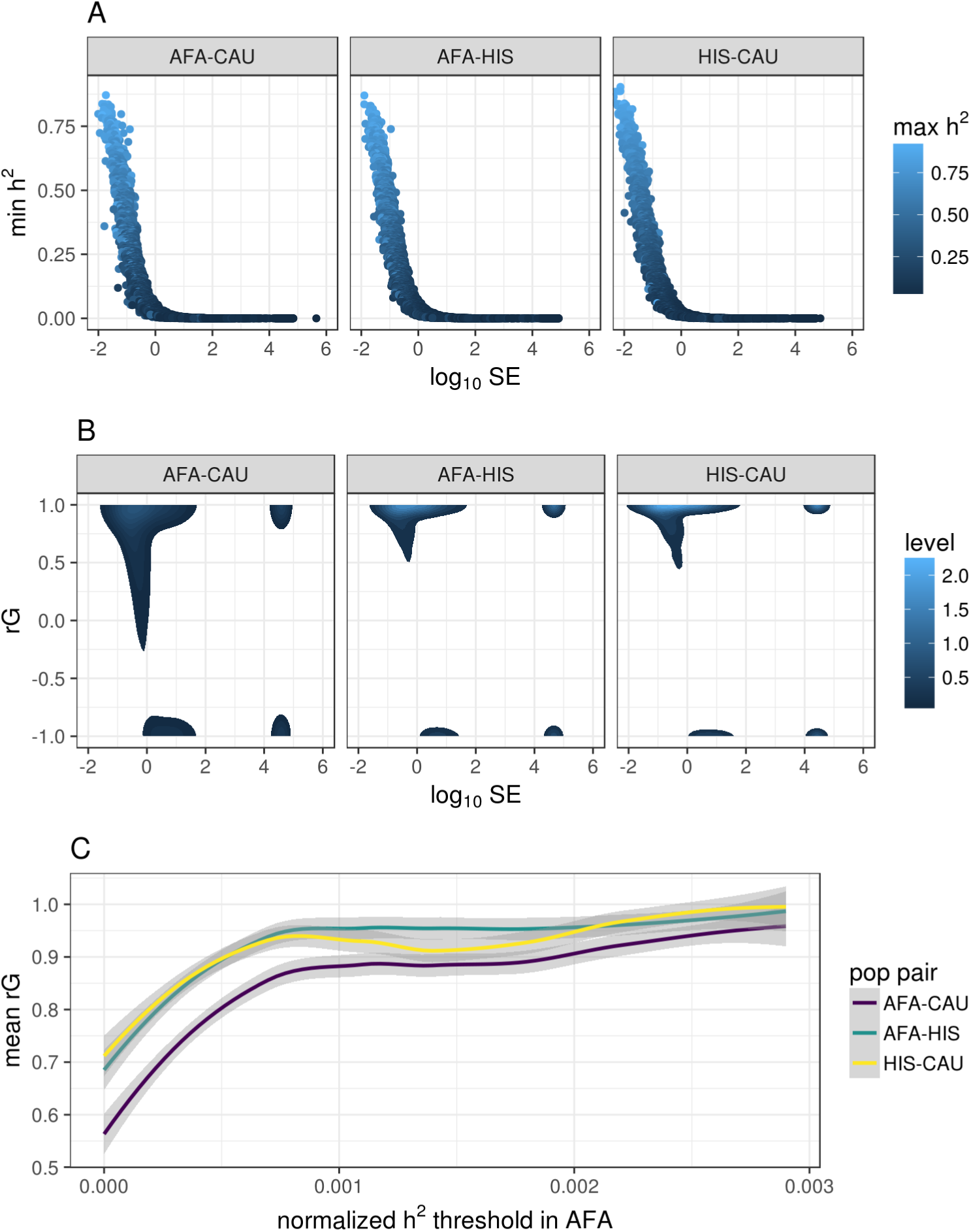
Higher heritability genes have higher genetic correlation (rG) and lower standard error (SE) estimates between MESA populations. (**A**) Pairwise population comparison of minimum heritability (h^2^) and rG standard (SE) for each gene. The y-axis is the minimum h^2^, the x-axis is the *log*_10_ SE of the rG estimate, and the points are colored according to the maximum h^2^ between the populations titling each plot. (**B**) rG compared to − *log*_10_ SE of the estimate. Genes with low SE are more likely to have a positive rG estimate. (**C**) Comparison of the genetic correlation between pairwise MESA populations and the subset of genes with normalized h^2^ greater than a given threshold in the AFA population. h^2^ estimates are normalized by the number of SNPs used in the estimate, i.e. those within 1Mb of each gene.

**S5 Fig.**
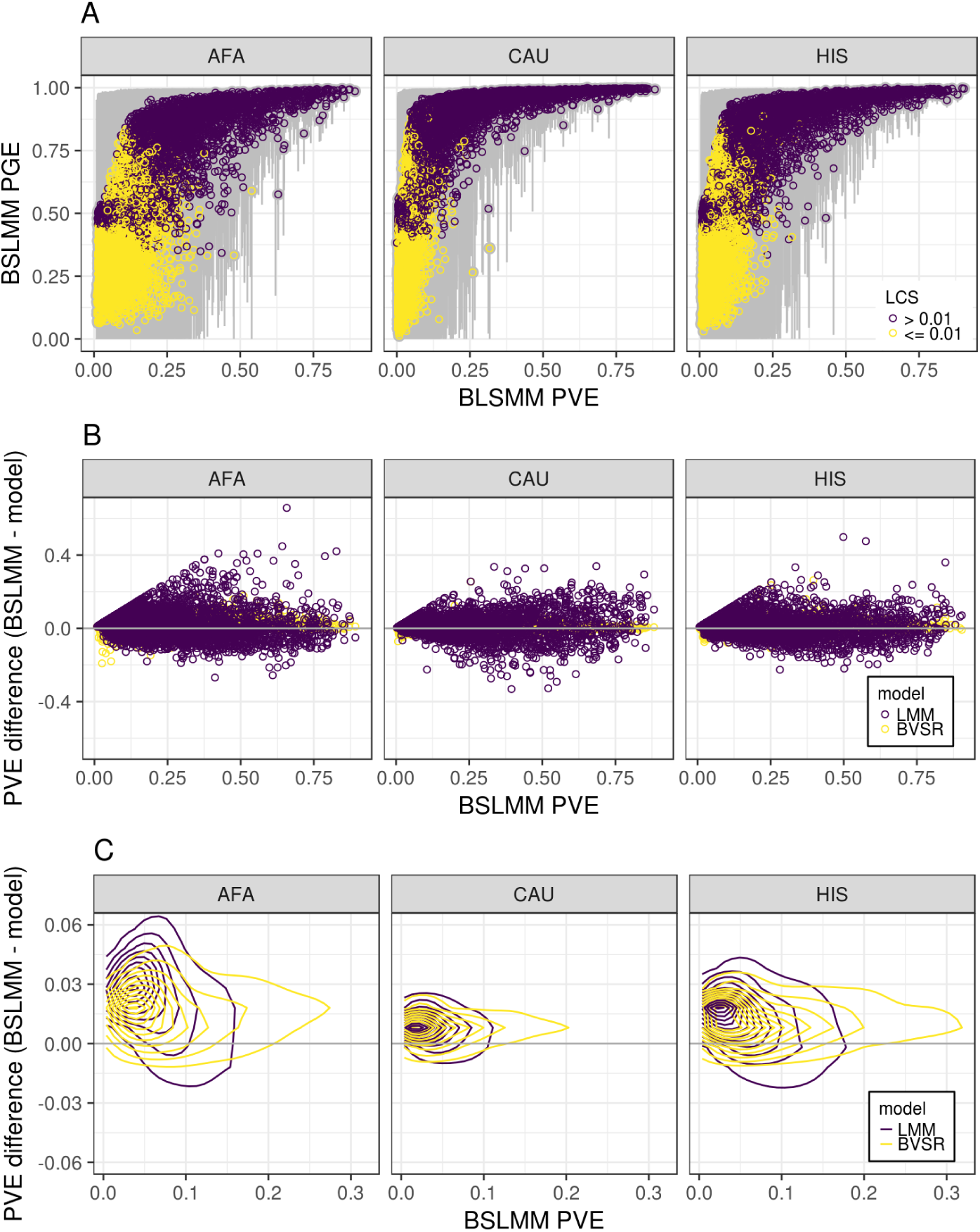
Comparison of gene expression proportion variance explained (PVE) estimates of models assuming different underlying genetic architectures. (**A**) Bayesian Sparse Linear Mixed Modeling (BSLMM) includes both sparse and polygenic components and estimates the total percent variance explained (PVE) and the parameter PGE, which represents the proportion of the genetic variance explained by sparse effects. The highly heritable genes (high PVE) have PGE near 1and therefore the local genetic architecture is sparse. There is not enough evidence to determine if the lower heritablility genes are more sparse or polygenic. (**B**) The difference between PVE of BSLMM and LMM or BVSR is compared to the BSLMM PVE across genes in MESA populations AFA, HIS, and CAU. (**C**) Zoomed in plot of **A** using contour lines from two-dimensional kernel density estimation to visualize where the points are concentrated. For both LMM and BVSR, the PVE difference values (y-axis) are above the horizontal line at zero indicating that both models perform worse than BLSMM. However, the difference between LMM and BSLMM is greater than between BVSR and BSLMM, which indicates sparse effects predominate for most genes.

**S6 Fig.**
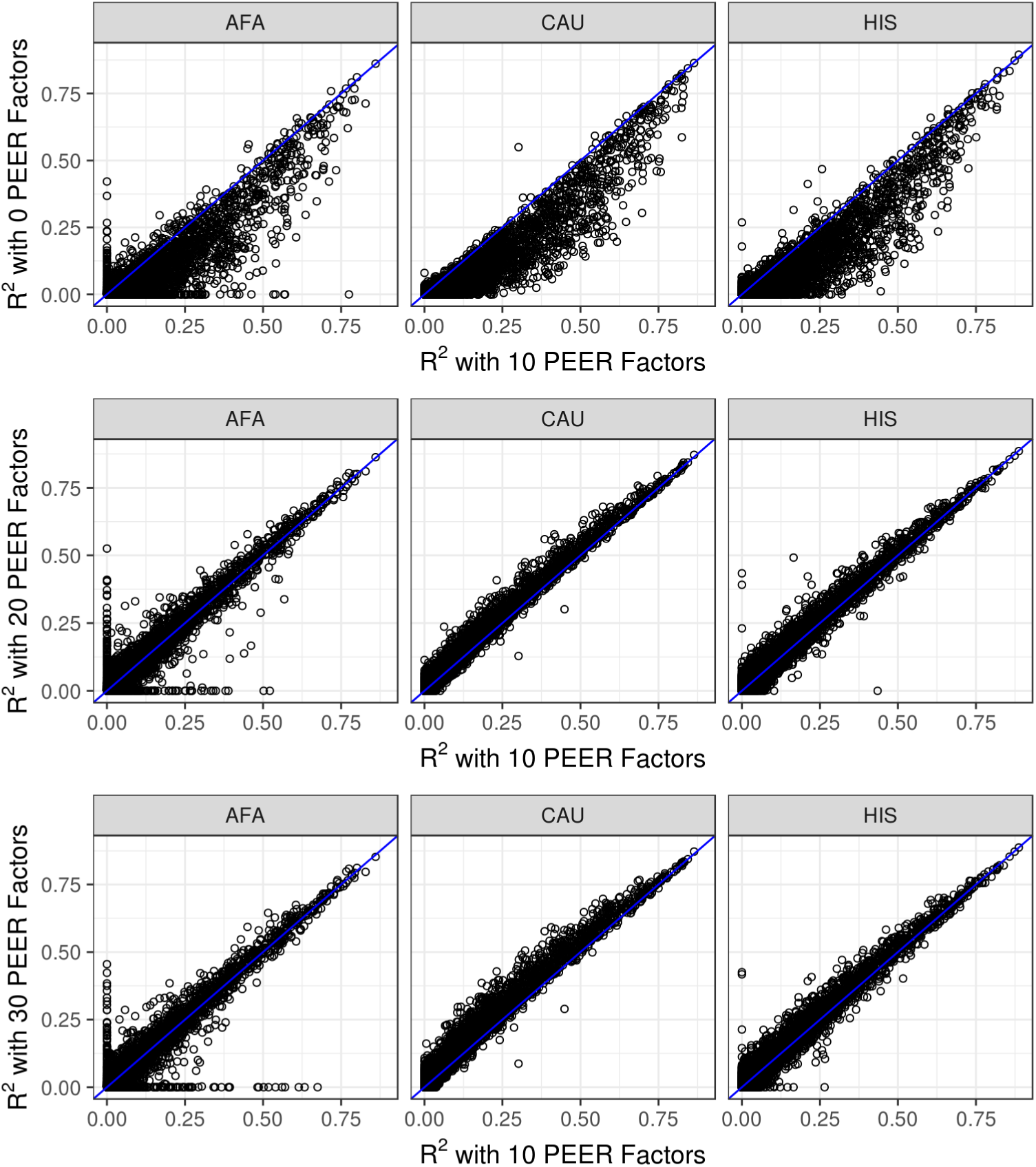
Consistent elastic net results with 10 or more PEER factors. Comparison of the elastic net (*α* = 0.5) cross-validated predictive performance R^2^ in models with different numbers of PEER factors as covariates. Across populations, models with 10 PEER factors shows increased predictive performance over 0 PEER factors, while models with 10, 20, or 30 PEER factors perform similarly.

**S7 Fig.**
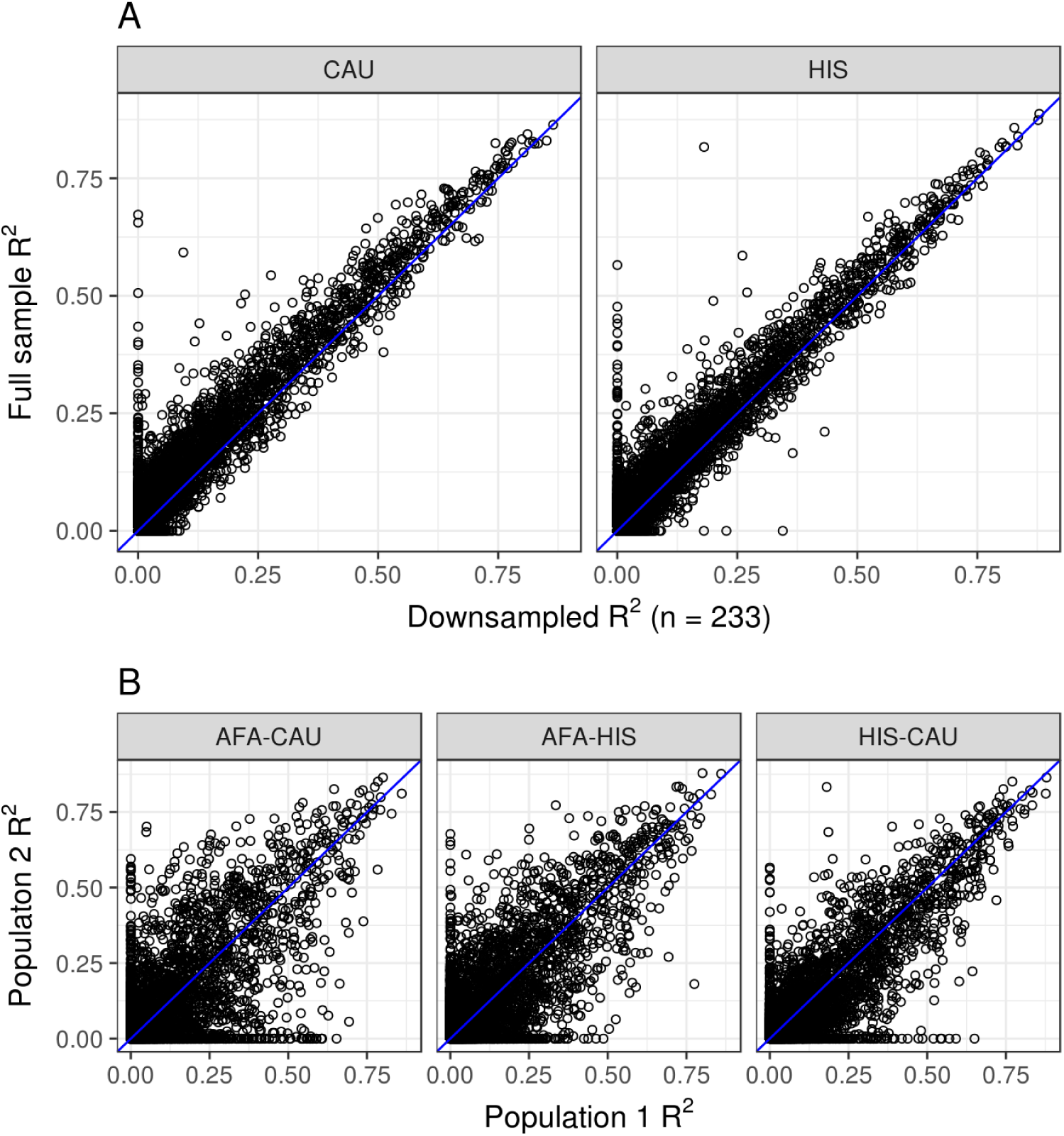
Consistent elastic net results with downsampled populations. The CAU and HIS populations were randomly downsampled to include the same sample size as AFA (n = 233). Predictive performance was measured within each population using nested cross-validation. (**A**) Comparison of the elastic net (*α* = 0.5) predictive performance R^2^ of the full sample to the downsampled population. Spearman correlations were 0.961and 0.966 for CAU and HIS sample comparisons, respectively. Comparison of predictive performance for each gene (R^2^) between each pair of populations. In each gray title box, population 1is listed first and population 2 is listed second. The identity line is shown in blue. The pairwise Spearman correlations (*ρ*) between genes are AFA-CAU downsample: *ρ* = 0.74, AFA-HIS downsample: *ρ* = 0.82, HIS downsample-CAU downsample: *ρ* = 0.87.

**S8 Fig.**
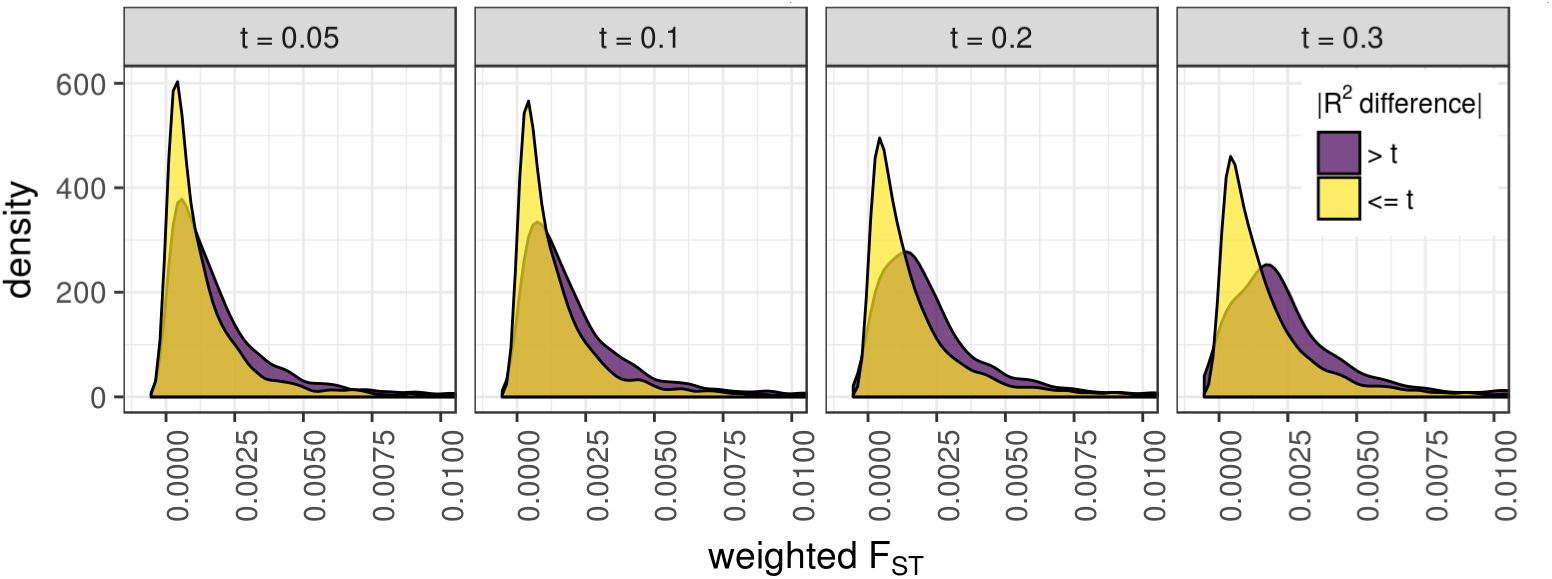
Fig Comparison of weighted F_ST_ between gene models with large (> t) and small (*<* = t) differences in predictive performance R^2^. For each gene model, weighted average F_ST_ was calculated by multiplying each beta from the elastic net model by that SNP’s F_ST_ before taking the mean across SNPs. The gene groups with the larger absolute value R^2^ difference between populations had significantly larger weighted F_ST_ at each difference threshold, t (Wilcoxon rank sum tests, *P <* 2.2 *×;* 10^−16^).

**S1 Table. Bonferroni significant S-PrediXcan results using gene expression prediction models from different MESA populations and summary statistics from a multiancestry GWAS of asthma.**

**S2 Table. All S-PrediXcan results using gene expression prediction models from different MESA populations and summary statistics from a multiancestry GWAS of asthma.**

